# Examining the influence of environmental factors on *Acanthamoeba castellanii* and *Pseudomonas aeruginosa* in co-culture

**DOI:** 10.1101/2024.01.17.575952

**Authors:** Rhiannon E. Cecil, Deborah R. Yoder-Himes

## Abstract

Exploration of interspecies interactions between microorganisms can have taxonomic, ecological, evolutionary, or medical applications. To better explore interactions between microorganisms it is important to establish the ideal conditions that ensure survival of all species involved. In this study, we sought to identify the ideal biotic and abiotic factors that would result in high co-culture viability of two interkingdom species, *Pseudomonas aeruginosa* and *Acanthamoeba castellanii*, two soil dwelling microbes. Long-term co-culture of these two organisms has traditionally been unsuccessful and usually results in high mortality for one or both organisms suggesting a predator-predator interaction may exist between them. In this study, we identified biotic and abiotic conditions that resulted in a high viability for both organisms in long-term co-culture, including optimizing temperature, nutrient concentration, choice of bacterial strains, and the initial ratio of interacting partners. These two species represent ideal partners for studying microbial interactions because amoebae act similarly to mammalian immune cells in many respects, and this can allow researchers to study host-pathogen interactions *in vitro*. Therefore, long-term interaction studies between these microbes might reveal the evolutionary steps that occur in bacteria when subjected to intense predation, like what occurs when pathogens enter the human body. The culture conditions characterized here resulted in high viability for both organisms for at least 14-days in co-culture suggesting that long-term experimental studies between these species can be achieved using these culture conditions.

## Introduction

*Pseudomonas aeruginosa* is an opportunistic human pathogen that is commonly isolated from other natural environments such as soil or freshwater bodies of water. It is a leading cause for nosocomial infections in the U.S. and can infect nearly every organ system in the mammalian body including the skeletal system, cardiovascular system, digestive system, integumentary system, reproductive system, and respiratory system [1–6]. Its ability to survive in such a wide range of habitats is largely due to its large, plastic genome leading to wide metabolic capabilities [7, 8]. *P. aeruginosa* is also known to reduce the size of its genome via gene deletions to become highly adapted to a specific niche within a human host [9, 10]. *P. aeruginosa* has a wide arsenal of virulence factors which gives it a competitive edge over both prokaryotic and eukaryotic competitors within natural and anthropogenic environments [11–24].

*Acanthamoeba castellanii* is a natural predator of *P. aeruginosa. A. castellanii* is a free-living, single-celled, lobose amoeba species and is a model organism to study phagocytosis [25, 26]. Amoebae and phagocytic human immune cells, such as macrophage, share many conserved proteins and traits such as P_21_-activated kinase, Ras and Rab proteins, PI phosphatases and P1 kinases, mannose binding proteins, and phagocytosis [27–30] Due to these similarities, it has been proposed that adaptations acquired by previously innocuous bacteria to escape/kill amoebae can, incidentally, result in virulence factors effective against human immune cells. One example organism for studying host-pathogen interactions between eukaryotes and facultative bacterial pathogens is *Legionella pneumophila,* which is an aquatic bacterium that survives and replicates intracellularly in eukaryotic cells. Amoebae are its natural host but, upon infecting humans, this organism is able to survive inside monocytes and macrophages using the same mechanisms of entry and intracellular survival strategies employed for their intracellular lifestyle within amoebae [31–33]. *A. castellanii* itself is also an opportunistic human pathogen with infections resulting in ocular keratitis and can also cause a rare form of encephalitis called *Acanthamoeba* granulomatous encephalitis [30]. These amoebae are found in both natural and anthropogenic water sources such as ponds or sink drains respectively. *A. castellanii* exists in two forms: a metabolically active, phagotrophic trophozoite form and a cryptobiotic cyst form. Its lifecycle involves a growth phase and two cellular differentiation stages: encystment and excystment [34]. In the metabolically active stage of the amoeba’s life cycle, the cell is referred to as a trophozoite. Trophozoites are motile, reproduce mitotically, and can phagocytose prey [35, 36]. Because *A. castellanii* is a proven predator of *P. aeruginosa* [37, 38] and can be found in the same types of environmental and man-made reservoirs, it would make for an ideal natural host to examine host-pathogen interactions.

One limitation to studying host-pathogen interactions using bacteria and amoeba is that often the interacting partners kill each other quite readily under certain abiotic and biotic conditions [37, 39–41]. This suggests that predator-predator interactions may also occur between these species rather than a strictly predator-prey interaction. Therefore, identifying conditions that maximize both the survival and the interaction between the partners is critical for understanding the molecular changes that are involved in the host-pathogen interactions and that may change over time.

In this study, we sought to optimize conditions to allow survival of *P. aeruginosa* and *A. castellanii* in co-culture for short- or long-term interaction experiments. We examined the effect of abiotic factors (temperature and concentration of nutrients) and biotic factors (bacterial isolate life history and concentration of interacting organisms) to maximize survival of both organisms in long-term co-culture. With optimal conditions identified, ecological and evolutionary interactions between these microorganisms can be further explored at the molecular and organismal level.

## Materials And Methods

### Strains used in this study, culturing, and cell maintenance

Strains used in this study are described in Table 1. *A. castellanii* strain 30010 was obtained from the ATCC. Clinical *P. aeruginosa* strains were obtained from Norton’s Children’s Hospital microbiology lab (strains B84725, B80398, B80427, B80422, B80425, B84723, and PA3) from cystic fibrosis sputum. Environmental *P. aeruginosa* strains were isolated from household sink drains in Louisville, Kentucky, U.S.A. (SRP 3151, SRP 17-047, and SRP 17-055) [42]. *A. castellanii* cells were maintained routinely in 15 mL of HL5 [25] in T-75 tissue culture flasks at room temperature (22°C) until they had reached confluence (∼ 5 days). The cells were passaged by using a cell scraper to dislodge the trophozoites, then a Pasteur pipet with suction was used to remove the culture from the flask, 1 mL of the culture was added back to the flask along with 14 mL of fresh HL5. *P. aeruginosa* cells were maintained in LB Lennox broth (with shaking) or on LB Lennox agar plates at 37°C unless otherwise noted.

**Table 1.** *Pseudomonas aeruginosa* strains used in this experiment and their respective phenotypes.

| Strain | Origin | Colony Phenotype | References |
| --- | --- | --- | --- |
| PA B80398 | Clinical – CF sputum sample | Normal | This study |
| PA B80427 | Clinical – CF sputum sample | Small colony | This study |
| PA B84725 | Clinical – CF sputum sample | Mucoid | This study |
| PA B84723 | Clinical – CF sputum sample | Normal | This study |
| PA B80422 | Clinical – CF sputum sample | Small colony | This study |
| PA B80425 | Clinical – CF sputum sample | Small colony | This study |
| PA 3 | Clinical – CF sputum sample | Normal | This study |
| SRP 3151 | Environmental – bathroom sink drain | Normal | [43-45] |
| SRP 17-047 | Environmental – bathroom sink drain | Normal | [43-45] |
| SRP 17-055 | Environmental – kitchen sink drain | Normal | [43-45] |

### General Method for Co-Culture Experiments

*A. castellanii* was cultured in 100% HL5 at room temperature in T-75 tissue culture flasks until confluent. Cells were scraped with a tissue culture scraper to dislodge cells from the surface and counted on a hemocytometer. Approximately 6x10^3^ cells of *A. castellanii* based on hemocytometer estimates, were added to 4 mL of fresh 100% HL5 in 6-well tissue culture-treated polystyrene plates. The cells were allowed to incubate in the HL5 for 30 minutes at room temperature to allow the cells to anneal to the well bottom. The HL5 was removed via a Pasteur pipet with vacuum suction and the cells were washed 2 times with 4 mL sterile PBS. Then 8 mL of fresh sterile indicated medium was added to each well. *A. castellanii* cells were cultured in monoculture or in co-culture with *P. aeruginosa* strains, grown to mid-log phase in 5 mL LB Lennox broth, under the conditions listed below. *A. castellanii* and *P. aeruginosa* cells were enumerated on the indicated days as described below.

### Co-Culture Variable Optimization

Strain optimization: Ten total P. aeruginosa strains were selected and are listed in Table 1. Each was cultured in LB Lennox broth and added to *A. castellanii* as indicated above. Co-cultures and mono-cultures were diluted into 8 mL sterile PBS in 6-well tissue culture plates for the duration of the interaction study to force bacterial-amoeabal interaction. Cultures were allowed to incubate over 7 days at 22°C and were enumerated at 3, 5, and 7 days post-inoculation.

Optimal medium concentration: *A. castellanii* and *P. aeruginosa strains* were cultured in monoculture or in co-culture in 6-well tissue culture plates containing 8 mL of 0.1% HL5, 1% HL5, 10% HL5, or 100% HL5 at 22°C for 14 days. Each strain was tested with biological triplicates. The cells were cultured in a 10 *A. castellanii* to 1 *P. aeruginosa* ratio. Every two days the supernatant was removed and replaced with 8 mL of the respective medium. *A. castellanii* and *P. aeruginosa* cells were enumerated on days 1, 7, and 14.

Optimal temperature: *A. castellanii* and *P. aeruginosa* strains were cultured in 8 mL of 1 % HL5 in 6-well tissue culture plates as described for the medium concentration experiment above except the cultures were incubated at either 22°C, 30°C, or 37°C in 1% HL5 medium for 14 days.

Ratio of *A. castellanii* and *P. aeruginosa*: *A. castellanii* and *P. aeruginosa* were cultured as describe for the medium concentration experiment above except the cultures were mixed at MOIs of 100:1, 10:1, 1:1, 1:10, or 1:100 amoebae:bacteria and cultures were grown at 22°C in 1% HL5 medium for 14 days.

### Enumeration of A. castellanii and P. aeruginosa

*A. castellanii* cells were enumerated via hemocytometer prior to the initiation of each experiment. *A. castellanii* was also subsequently enumerated via bright field microscopy at 400X magnification by examining three replicate images within each well to assess the number of surviving trophozoites and cysts present during and at the end of each experiment. *P. aeruginosa* cells were enumerated via serial dilution and plating on Lennox agar.

### Statistics

GraphPad Prism v5.04 was used for data analysis. Data were analyzed with either t-tests, one-way ANOVAs with either Dunnett’s or Tukey’s post-test, or two-way ANOVAs as indicated in each analysis. The effects of life history could not be directly analyzed in Figs 3 – 5 due to lack of statistical power as only one environmental strain was tested in those experiments.

## Results

To better understand the relational dynamics between *P. aeruginosa* and *A. castellanii* in co-culture, we optimized several variables to ensure long-term survival of both partners. We sought to understand how life history (clinical versus environmental) of the bacterial strains impacted the relationship with the amoebae. Further, we sought to optimize abiotic factors (medium and temperature) of the cultures, and biotic factors (ratio of bacteria to amoebae) in order to extend co-culture survival of both organisms. The contribution of each partner to survival was also explored as *P. aeruginosa* serves as a food source for the amoebae and it is possible that amoebae excretions/secretions or the amoebae themselves could serve as a food source for *P. aeruginosa* in co-culture with low nutrient growth medium.

### Impact of P. aeruginosa strain on A. castellanii survival

The ecological niche from which the *P. aeruginosa* strains originated may play a strong role in their interaction with *A. castellanii* as different phenotypes of *P. aeruginosa* display different virulence characteristics. For example, mucoid colony phenotypes or small colony variants of *P. aeruginosa* are associated with chronic infections and lower virulence factor expression while normal phenotypes are associated with acute infections and higher virulence factor expression [46–48]. Ten different *P. aeruginosa* strains (Table 1) were tested for their effects on co-culture survival dynamics with *A. castellanii* over 7 days to determine if any of the *P. aeruginosa* strains were either killed by *A. castellanii* or conversely killed *A. castellanii*. Further, we determined whether *A. castellanii* cells remained in their actively growing state (trophozoite) or encysted in the presence of these strains.

All *P. aeruginosa* strains were able to survive in the presence of *A. castellanii* over the course of the experiment (Fig 1A). After only 24 hours, several of the clinical strains, including B80427, B80425, and PA3, showed a small reduction in survival compared to all of the strains grown in the absence of amoebae; however, all of them rebounded by day 3 or 5. Interestingly, PA B80425 did not remain at elevated levels on day 7 and instead showed another reduction in growth compared to mono-culture suggesting that it is less robust than the other strains when in the presence of amoebae. The environmental isolates all survived at a greater concentrations in co-culture than in mono-culture over all 4 time points suggesting that these particular strains, at least, were well-prepared for co-culture with amoebae which is not surprising as both *P. aeruginosa* and *A. castellanii* are likely to share environmental niches and *A. castellanii* has been shown to aid in the extracellular survival of other bacterial species [49, 50]. Raw survival counts for *P. aeruginosa* are provided in S1 Fig.

**Fig 1.**
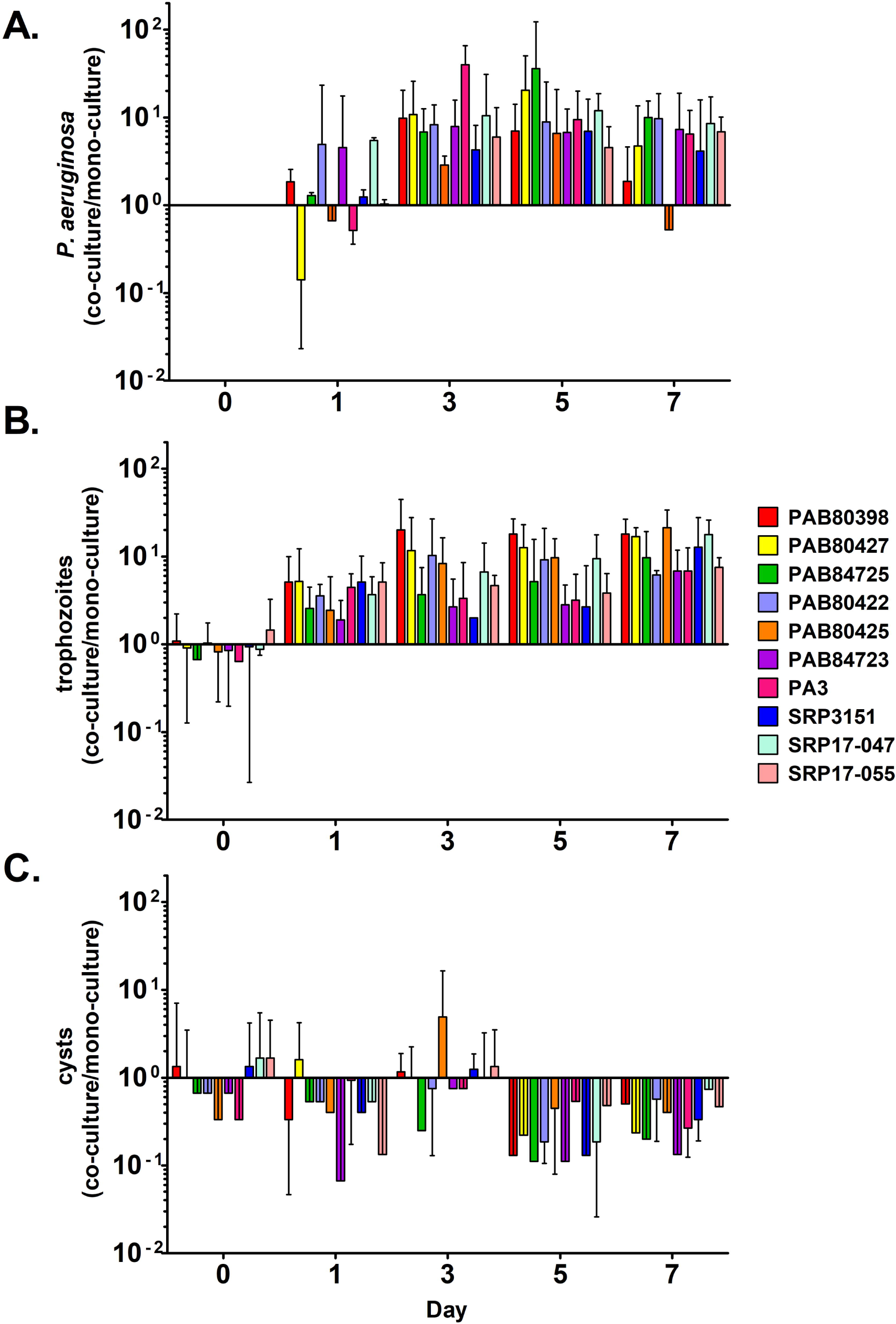
*A. castellanii* exhibits increased trophozoite survival and decreased encystment in co-culture with *P. aeruginosa* compared to monoculture. In all panels, 10^0^ represents the number of cells or CFUs present in monoculture at each time point. (A) Ratio of *P. aeruginosa* concentrations in co-culture with *A. castellanii* to *P. aeruginosa* survival in monoculture. (B) Ratio of the number of *A. castellanii* trophozoites present in co-culture with each *P. aeruginosa* strain compared to monoculture. (C) Number of *A. castellanii* cysts present in co-culture with *P. aeruginosa* compared to monoculture. Data analyzed with two-way ANOVA (S1 Table).

*A. castellanii* survived and stayed in trophozoite form when in co-culture with all 10 *P. aeruginosa* strains at days 1, 3, 5, and 7 compared to monoculture (Fig 1B), perhaps owing to the food source of *P. aeruginosa* in these cultures. *A. castellanii* trophozoite numbers were generally highest when co-cultured with PA B80389 (normal colony, clinical isolate), PA B80427 (small colony, clinical isolate), PA B80425 (normal colony, clinical isolate), and SRP 17-047 (normal colony, environmental isolate) (Fig 1B). Raw *A. castellanii* trophozoite and cyst counts are provided in S2 Fig. Data tables for 2-way ANOVA results are available in S1 table.

Encystment generally decreased over the course of the experiment though this effect was variable depending on the *P. aeruginosa* strain with which *A. castellanii* was co-cultured (Fig 1C). This variability is most likely due to the low numbers of cysts overall in most conditions except in the monoculture in which encystment occurred to a much higher concentration than any of the co-culture conditions (S2 Fig).

From this experiment, it appears that *A. castellanii* was able to survive with all 10 *P. aeruginosa* strains tested. Trophozoite survival was greater in co-culture with *P. aeruginosa* than in monoculture indicating that *A. castellanii* is able to readily predate on all of these *P. aeruginosa* isolates. We also concluded that *P. aeruginosa* grew more robustly in co-culture with *A. castellanii* than in monoculture which suggests that the amoebae are either serving as a food source themselves or are producing products that are able to be metabolized by *P. aeruginosa*.

To assess whether life history of the bacterial isolates is important for amoebae survival, we grouped the strains, those from humans and those from sink drains, and tested for significant differences in the number of surviving trophozoites. On average, *A. castellanii* trophozoites were more abundant when co-cultured with clinical *P. aeruginosa* strains than when cultured with environmental strains (Fig 2). These results were not significantly different, suggesting that origin of isolation does not appear to influence these interactions. However, we should note that these drain isolates may have recently been associated with human hosts as they were isolated from households, and this could mean that they are more human-adapted rather than true environmental (i.e. non-human associated).

**Fig 2.**
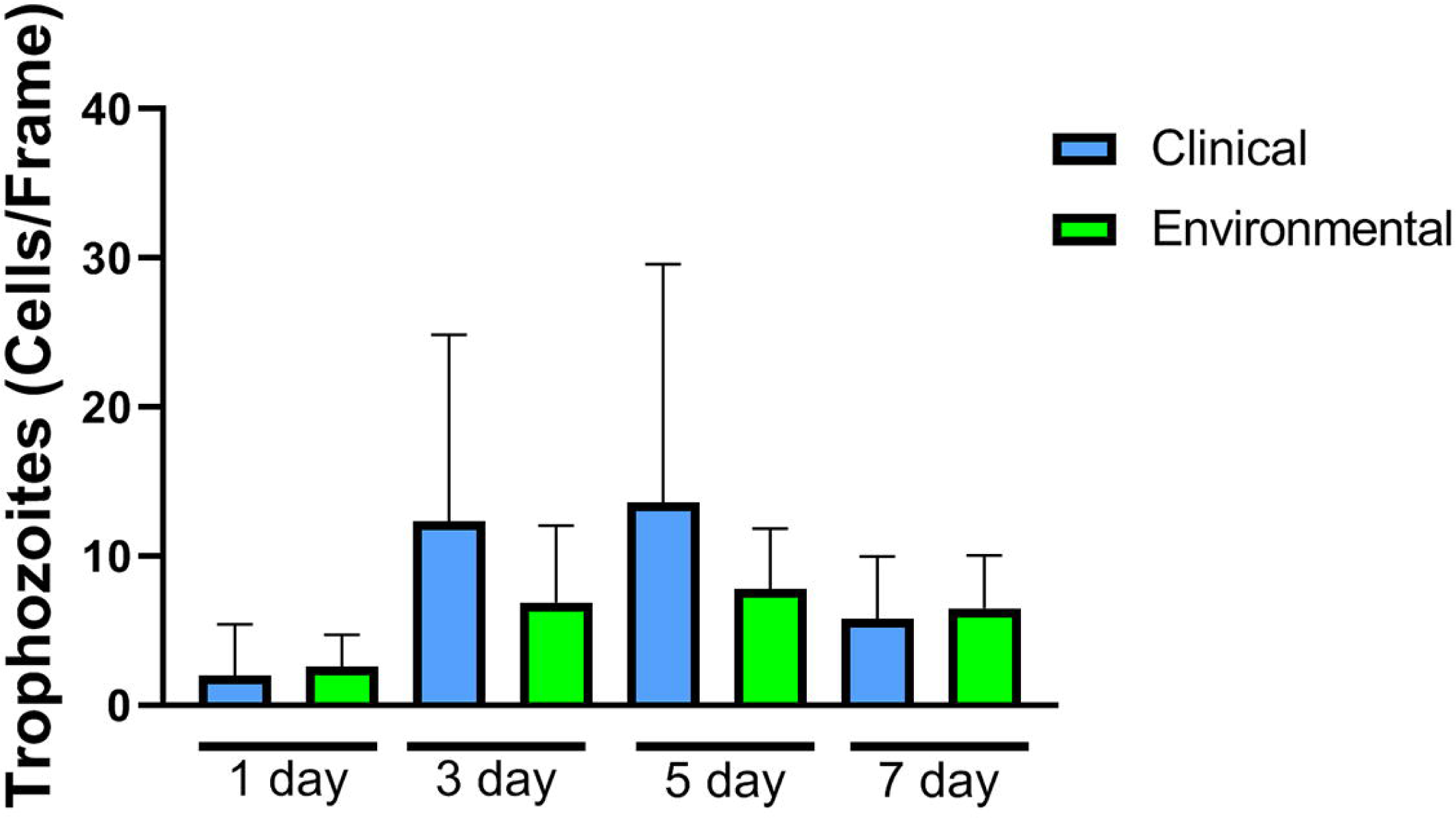
*A. castellanii* survival in co-culture with clinical versus environmental *P. aeruginosa* strains. The mean survival of *A. castellanii* trophozoites cultured with the 7 clinical isolates or 3 environmental isolates are shown. Data analyzed with t-test between groups at each time point. There were no significant differences between the two groups at any time point.

Four of the ten *P. aeruginosa* strains tested were chosen for further testing. Three strains were of clinical origin (B80398, B84725, and B80427) and represent three major colony phenotypes (normal, mucoid, and small colony variant respectively). The other strain was of environmental origin, SRP 3151 (normal phenotype). The three clinical isolates were chosen based on their colony phenotypes and the environmental isolate was chosen due to its use in other studies [43–45] and because its genome has been sequenced, data unpublished.

### Influence of nutrient concentration

Nutrient levels could strongly influence the interactions between amoebae and bacteria. *A. castellanii* grows axenically in nutrient rich media, such as HL5, but can also grow in nutrient poor media using bacteria as a food source. To better understand how nutrient levels influenced co-culture survival for *P. aeruginosa* and *A. castellanii*, *A. castellanii* cells were cultured in monoculture or in co-culture with *P. aeruginosa* in 0.1% HL5, 1% HL5, 10% HL5, and 100% HL5 for up to 14 days. We hypothesized that the low nutrient conditions (0.1% or 1% HL5) would promote survival of *A. castellanii* in co-culture with *P. aeruginosa* as co-culturing these organisms in high nutrient medium results in high *A. castellanii* mortality based on pilot experiments.

*P. aeruginosa* survived at high concentrations (between 10^6^ and 10^10^ CFU/mL) in monoculture and in co-culture with *A. castellanii* in all nutrient concentrations tested (Fig 3 A - C). However, *P. aeruginosa* CFU/mL tended to be highest when cultured in 100% HL5 compared to the other concentrations tested particularly at days 7 and 14 (Fig 3B and C). *A. castellanii* trophozoites in monoculture were significantly more abundant when cultured in 100% HL5 compared to the all of the lower concentrations of HL5 tested (Fig 3 D-E). Amoebae trophozoite numbers co-cultured with 3 of the 4 *P. aeruginosa* strains were not significantly different between the nutrient concentration groups after one day (Fig 3D). In contrast, *A. castellanii* trophozoites were significantly more abundant when co-cultured in 1% HL5 than in 0.1%, 10%, or 100% HL5 at days 7 and 14 (Fig 3E and C). Taken together, we can conclude that 1% HL5 is the optimal concentration of HL5 for co-culture of *A. castellanii* and *P. aeruginosa* for long-duration experiments; however, this concentration of HL5 does not support the growth/survival of *A. castellanii* in monoculture.

**Fig 3.**
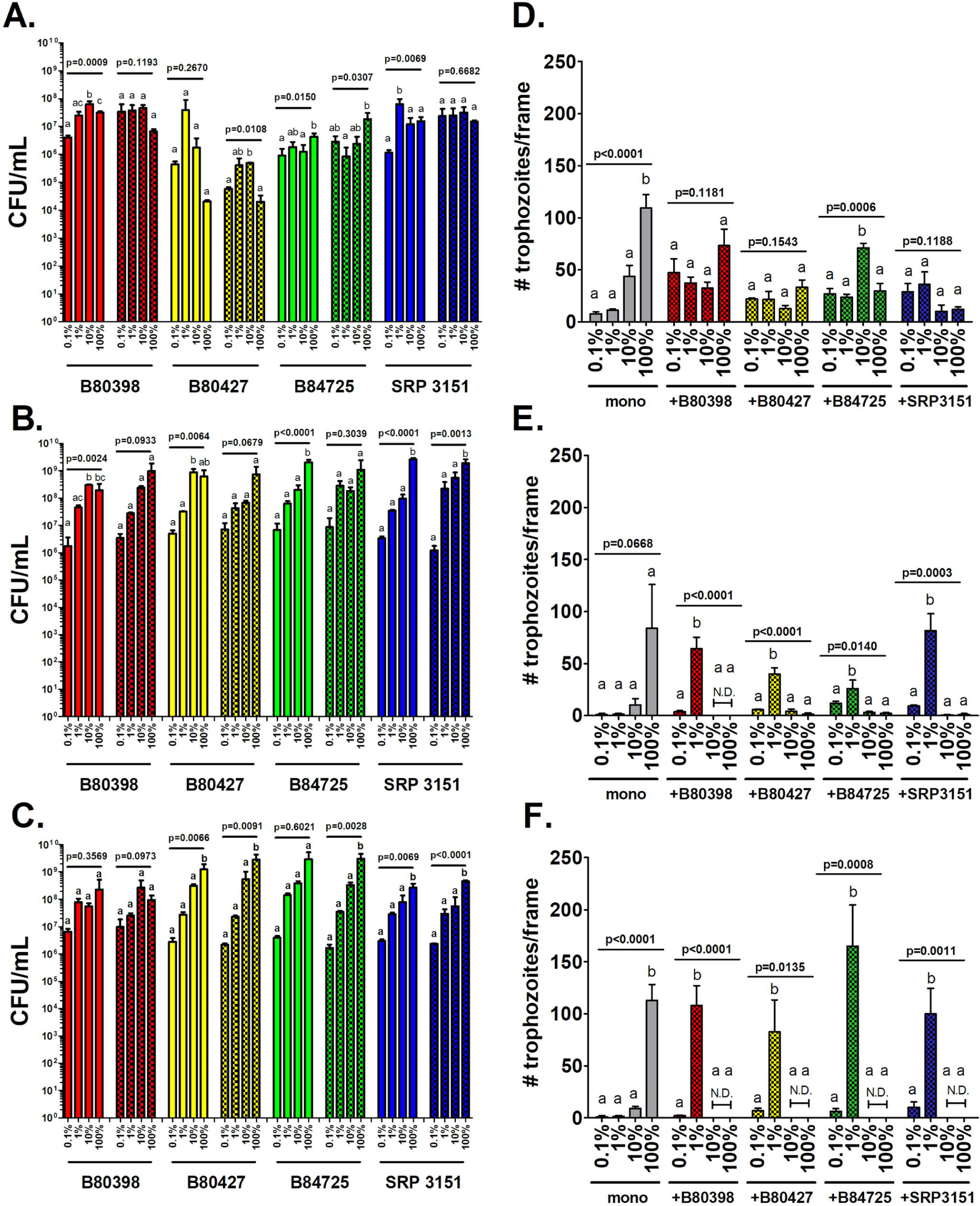
*A. castellanii* trophozoite survival in co-culture with *P. aeruginosa* is maximized when cultured in 1% HL5. *A. castellanii* was cultured in monoculture or in co-culture with four *P. aeruginosa* strains in 0.1% HL5, 1% HL5, 10% HL5 and 100% HL5 at 22°C. *P. aeruginosa* cells and *A. castellanii* trophozoites were enumerated on days (A) 1, (B) 7, and (C) 14. Data were analyzed with one-way ANOVA with Tukey’s post-test (n = 3 biological replicates). Lowercase letters indicate individual comparisons within each strain (but not between strains) and overall p-values for each group are indicated for each independent group comparison.

### Effect of temperature on co-culture population dynamics

Temperature is another abiotic factor that could have an immense effect on the co-culture survival of one or both of the studies organisms. It has been shown that *P. aeruginosa* alters its virulence factor expression depending on culture temperatures [51, 52]; therefore, it is imperative to determine how temperature influences the interactions between these pathogens. We hypothesized that room temperature (22°C) would be the optimal growth temperature for survival of both partners when co-culturing these two organisms as 22°C is roughly the temperature these organisms would experience in sink drains. The optimal growth temperature for *A. castellanii* ranges from 22°C to 32°C [53] . *P. aeruginosa* is able to survive/grow at a temperature range from 4°C to 42°C, with 37°C being its optimal growth temperature [51].

For this experiment *A. castellanii* was cultured in monoculture and in co-culture with *P. aeruginosa* for 14 days in 1% HL5 at 22°C, 30°C, and 37°C. Cultures were examined at three time points: 1-, 7-, and 14-days post-inoculation. In general, *P. aeruginosa* grew to higher concentrations at 37°C compared to 30°C or 22°C as expected, though growth was robust 7- and 14-days post-inoculation at all temperatures for all strains (Fig 4A – C). For *A. castellanii*, the number of trophozoites in monoculture was relatively high at day 1 but all cells had encysted or died 7- and 14-days post-inoculation (Fig 4D – F). This was expected as 1% HL5 is nutrient poor and does not support metabolically active trophozoites resulting in the conversion to metabolically dormant cysts in monoculture. In co-culture, there were usually more *A. castellanii* trophozoites present in the 22°C condition than were present in the 30°C or 37°C conditions at all three time points (Fig 4D – F). One exception to this is the high abundance of *A. castellanii* observed when co-cultured with SRP3151 at day 7 at 37°C (Fig 4E). However, because this trend was not observed at day 14, this is likely due to biological variability. Taken together, these results indicate that 22°C is a suitable temperature to maintain the survival of *P. aeruginosa* with the vegetative form of *A. castellanii* in co-culture.

**Fig 4.**
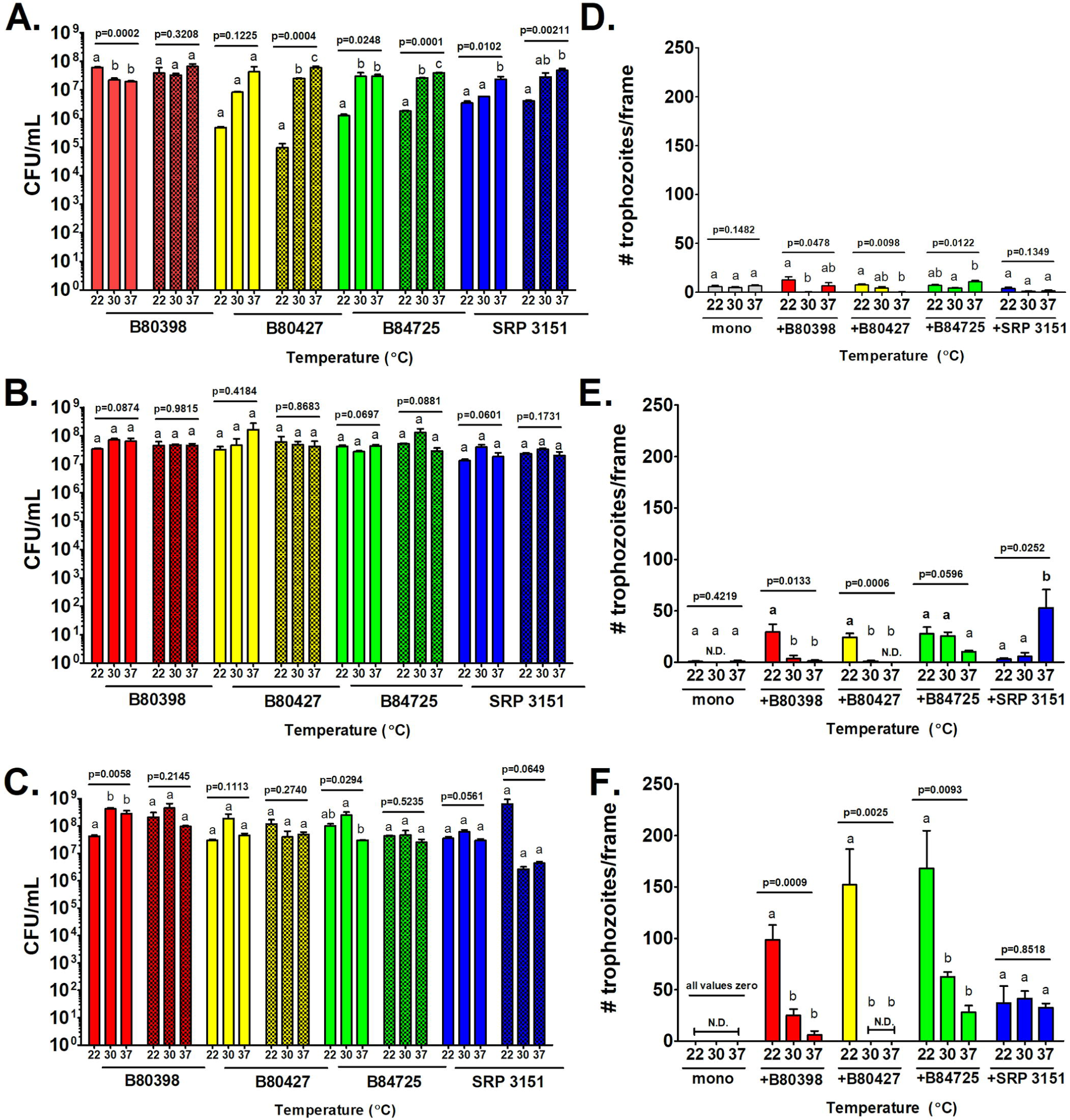
22°C is the ideal temperature for *A. castellanii* co-cultured with *P. aeruginosa*. *A. castellanii* was cultured in monoculture and in co-culture with 4 *P. aeruginosa* strains in 1% HL5 at 22°C, 30°C, or 37°C for 14 days. *A. castellanii* trophozoites and *P. aeruginosa* were enumerated via direct cell count at 400X magnification at (A and D) 1 day, (B and E) 7 days, and (C and F) 14 days. Panels A, B, and C represent *P. aeruginosa* survival while panels D, E, and F represent number of *A. castellanii* trophozoites. Data were analyzed with one-way ANOVA with Tukey’s post-test (n = 3 biological replicates). Lowercase letters indicate individual comparisons within each strain (but not between strains) and overall p-values for each group are indicated for each independent group comparison.

### The influence of inoculation ratios on co-culture dynamics

Since *A. castellanii* is an avid predator of *P. aeruginosa,* it is possible that a high amoebae:bacterium inoculation ratio will result in the eradication of the bacteria. It is also known that at a low amoebae:bacterium ratios, *P. aeruginosa* rapidly kills *A. castellanii* [37]. Therefore, we asked whether there is a middle ground that allows for the survival of both partners. We hypothesized that a starting concentration of less *P. aeruginosa* than *A. castellanii* would result in higher *A. castellanii* trophozoite viability in co-culture while not having a large impact on *P. aeruginosa* survival as *P. aeruginosa* has a much shorter doubling time than the amoebae.

To test this hypothesis, *A. castellanii* and *P. aeruginosa* were co-cultured with different starting ratios of *A. castellanii* to *P. aeruginosa* to identify which inoculation ratio resulted in the highest viability of *A. castellanii* trophozoites and *P. aeruginosa* cells over time. The cells were combined in ratios of 100:1, 10:1, 1:1, 1:10, or 1:100 amoebae:bacteria. The cells were incubated at 22°C and enumerated at 1, 7, and 14 days. On days 1 and 7, *P. aeruginosa* SRP3151 allowed for the most robust amoebae retention compared to the other *P. aeruginosa* strains on days 1 and 7 but this effect was not observed by day 14 when co-cultures with *P. aeruginosa* B80398, B84725, and SRP3151 had similar levels of trophozoites (Fig 5). *P. aeruginosa* B80427, the small colony variant supported the least amount of trophozoites at day 14 overall. We also observed that amoebae survival and proliferation in co-culture at these different ratios varied depending on *P. aeruginosa* strain. Monocultures of *A. castellanii* and *P. aeruginosa* were not assessed in this experiment as ratios of 1:0 or 0:1 amoebae:bacteria would not further our understanding of co-culture dynamics; furthermore, we have already established the growth patterns of each of these species in % HL5 at room temperature in the previous experiments mentioned in this study.

**Figure 5.**
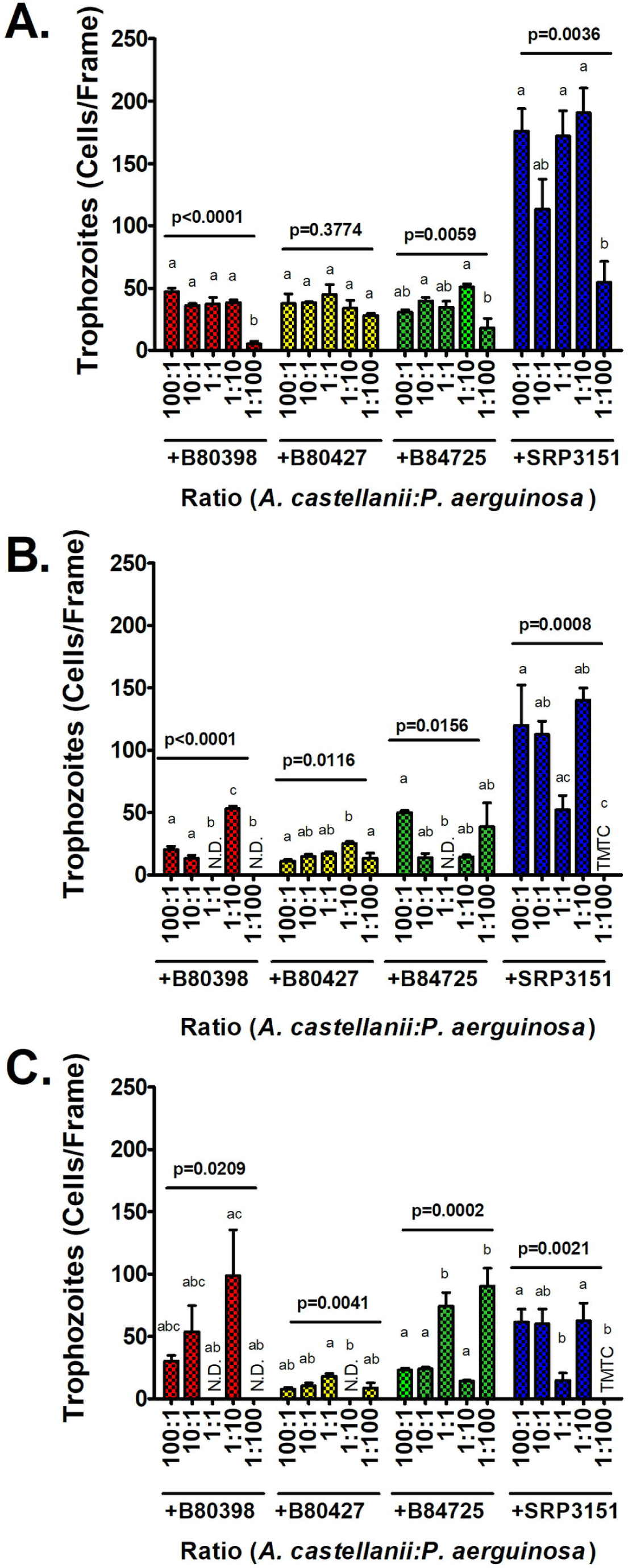
An initial dose of 10 *A. castellanii* to 1 *P. aeruginosa* cell results in the highest viable trophozoite yield overall. *A. castellanii* (AC) was cultured in monoculture and in co-culture with 4 *P. aeruginosa* (PA) strains at ratios of 100 AC to 1 PA, 10 AC to 1 PA, 1 AC to 1 PA, 1 AC to 10 PA, and 1 AC to 100 PA in 1% HL5 at 22°C for 14 days. *A. castellanii* trophozoites were enumerated via direct cell count at 400X magnification at (A) 1 day, (B) 7 days, and (C) 14 days. Data analyzed with one-way ANOVA with Tukey’s post-test (n = 3 biological replicates). Lowercase letters indicate individual comparisons within each strain (but not between strains) and overall p-values for each group are indicated for each independent group comparison.

## Discussion

*A. castellanii* and *P. aeruginosa* can be cultured together with high viability of both partners if the correct abiotic conditions are utilized. Low nutrient culture medium (1% HL5) is imperative to the co-culture survival of both organisms as high concentrations of nutrients resulted in the death of *A. castellanii* in co-culture with *P. aeruginosa*. Similarly, temperature is also an important abiotic factor as 22°C resulted in the most robust survival of *A. castellanii* when co-cultured with multiple *P. aeruginosa* strains while 37°C resulted in a significant reduction in viable *A. castellanii* cells in co-culture. Biotic factors such as life history of *P. aeruginosa* and the initial inoculum dose of each organism proved to have strain-dependent impacts on *A. castellanii* survival in co-culture.

Our results with respect to nutrient requirements are consistent with other bacteria-*A. castellanii* co-culture systems and demonstrate that *A. castellanii* is better equipped to tolerate co-culture with many bacterial species, including *P. aeruginosa,* under low nutrient conditions compared to high nutrient conditions as well as below a 100:1 bacteria to amoebae ratio [37]. Our experiments revealed that 22°C is a good co-culture temperature for these two species while 37°C resulted in high morbidity for *A. castellanii* when co-cultured with most of our *P. aeruginosa* strains tested. This result is similar to previously published literature that described experiments between *A. castellanii* and *Yersinia enterocolitica* where *A. castellanii* viability was reduced in both monoculture and co-culture at 37°C but both *Y. enterocolitica* and *A. castellanii* remained viable in the lower temperatures tested (7°C and 25°C) [54]. Similar results were also observed between *A. castellanii* and *Campylobacter jejuni* where *A. castellanii* viability was inhibited by the *C. jejuni* at 37°C but not at 25°C [55].

Increased survival of *A. castellanii* co-cultured with *P. aeruginosa* in lower-nutrient culture media and at temperatures below 37°C could be due to the fact that expression of virulence factors by *P. aeruginosa* is metabolically costly and some virulence factors have been shown to only be expressed at 37°C in clinical isolates [52]. Though it could be argued that long-term experimental evolution systems utilizing *P. aeruginosa* should be conducted at 37°C because it is the temperature at which *P. aeruginosa* is able to express many of its virulence factors and is also the temperature these organisms would experience inside a human host. It can also be argued that 22°C is the temperature these organisms would experience in their natural, non-mammalian habitat and thus would be the better temperature to assess co-culture survival and co-evolution between them.

Life history and colony phenotype did not have a substantial impact on *A.castellanii* survival in co-culture at an initial dose of 1 amoeba to 1 bacterial cell; however, there is a marked impact on *A. castellanii* survival when the initial dose of amoebae to bacteria is altered. *P. aeruginosa* strains from clinical origin proved to be more deadly overall to the amoebae than the environmental isolate tested by 14-days of co-culture in most of the ratios tested. Interestingly, this result is opposite to results observed when *A. castellanii* is co-cultured with clinical and environmental strains of *L. pneumophila,* though the only dose tested in that study was 1 amoebae to 10 bacteria cells [56]. Our results could be due to the fact that some of these clinical isolates could have been recently acquired by the host and thus their life history would be more liken to the environmental isolates tested; however, this is unlikely, particularly with respect to the small colony and mucoid isolates as these phenotypes (*P. aeruginosa* B84725 and B80427 respectively) are associated with chronic infection and heretofore are rarely if at all isolated from non-host-associated systems. Conversely, it is also possible that the environmental isolates could have recently been excreted from a human host into the bathroom/kitchen sink drains from which they were isolated, and we have no means to rule out this possibility.

The results of these experiments give insights into the complicated dynamics of polymicrobial communities when multiple species can act as predators of other community members, which is a common trope in the microbial world but less so in the animal kingdom where the trophic levels are more defined. Researchers can also use this information to look further into these specific interactions. For example, with the co-culture variables identified, it is possible to increase the duration of co-culture of *A. castellanii* and *P. aeruginosa* without needing to replenish the amoebae or bacteria for long-term experimental evolution experiments. This might be useful for studies that seek to test broad fundamental evolutionary or ecological hypotheses in addition to further understanding other polymicrobial, interkingdom interactions.

## Supporting Information

**S1 Fig. *Pseudomonas aeruginosa* viable cell count in monoculture and in co-culture with *Acanthamoeba castellanii* in PBS for 7-days.** Viable cell counts of *P. aeruginosa* strains are provided for monoculture (solid bars) and co-culture with *A. castellanii* (patterned bars). (A) B80398, (B) B80427, (C) B84725, (D) B80422, (E) PA B80425, (F) PA B84723, (G) PA3, (H) SRP 3151, (I) SRP 17-047, and (J) SRP 17-055. Data analyzed with t-tests comparing monoculture to co-culture at each time point.

**S2 Fig. *Acanthamoeba castellanii* cell count in monoculture and in co-culture with *Pseudomonas aeruginosa* in PBS for 7-days.** Direct cell counts of *A. castellanii* (A) trophozoites and (B) cysts in monoculture and in co-culture with *P. aeruginosa* are provided. Data analyzed with 1-way ANOVA with Dunnett’s post-test comparing each co-culture condition to monoculture at each time point.

**S1 Table. Two-way ANOVA on the effects of *P. aeruginosa* strain and time on the survival of *P. aeruginosa* and *A. castellanii* in PBS.** Table summarizes the 2-way ANOVA analysis output for figure 1.

*DF (degrees of freedom), SS (summary of squares), MS (mean of squares)

## References

1. Cerioli M, Batailler C, Conrad A, Roux S, Perpoint T, Becker A, et al. *Pseudomonas aeruginosa* implant-associated bone and joint infections: Experience in a regional reference center in France. Frontiers in Medicine. 2020;7:513242.

2. Bouchart F, Dubar A, Bessou JP, Redonnet M, Berland J, Mouton-Schleifer D, et al. *Pseudomonas aeruginosa* coronary stent infection. The Annals of thoracic surgery. 1997;64(6):1810–3.

3. Markou P, Apidianakis Y. Pathogenesis of intestinal *Pseudomonas aeruginosa* infection in patients with cancer. Frontiers in cellular and infection microbiology. 2014;3:115.

4. Spernovasilis N, Psichogiou M, Poulakou G. Skin manifestations of *Pseudomonas aeruginosa* infections. Current Opinion in Infectious Diseases. 2021;34(2):72–9.

5. Rana K, Thaper D, Vander H, Prabha V. *Pseudomonas aeruginosa*: A risk factor for fertility in male mice. Reproductive Biology. 2018;18(4):450–5.

6. Mendes OR. The challenge of pulmonary *Pseudomonas aeruginosa* infection: How to bridge research and clinical pathology. Viral, Parasitic, Bacterial, and Fungal Infections: Elsevier; 2023. p. 591–608.

7. Stover CK, Pham XQ, Erwin A, Mizoguchi S, Warrener P, Hickey M, et al. Complete genome sequence of *Pseudomonas aeruginosa* PAO1, an opportunistic pathogen. Nature. 2000;406(6799):959–64.

8. Winsor GL, Griffiths EJ, Lo R, Dhillon BK, Shay JA, Brinkman FS. Enhanced annotations and features for comparing thousands of *Pseudomonas* genomes in the *Pseudomonas* genome database. Nucleic acids research. 2016;44(D1):D646–D53.

9. Smith EE, Buckley DG, Wu Z, Saenphimmachak C, Hoffman LR, D’Argenio DA, et al. Genetic adaptation by *Pseudomonas aeruginosa* to the airways of cystic fibrosis patients. Proceedings of the National Academy of Sciences of the United States of America. 2006;103(22):8487–92. Epub 2006/05/12. doi: 10.1073/pnas.0602138103. PubMed PMID: 16687478; PubMed Central PMCID: PMCPMC1482519.

10. Lucchetti-Miganeh C, Redelberger D, Chambonnier G, Rechenmann F, Elsen S, Bordi C, et al. *Pseudomonas aeruginosa* Genome Evolution in Patients and under the Hospital Environment. Pathogens (Basel, Switzerland). 2014;3(2):309–40. Epub 2014/12/02. doi: 10.3390/pathogens3020309. PubMed PMID: 25437802; PubMed Central PMCID: PMCPMC4243448.

11. Hauser AR. The type III secretion system of *Pseudomonas aeruginosa*: infection by injection. Nature Reviews Microbiology. 2009;7(9):654–65.

12. Horna G, Ruiz J. Type 3 secretion system of *Pseudomonas aeruginosa*. Microbiological Research. 2021;246:126719.

13. Britigan BE, Roeder TL, Rasmussen GT, Shasby DM, McCormick ML, Cox CD. Interaction of the Pseudomonas aeruginosa secretory products pyocyanin and pyochelin generates hydroxyl radical and causes synergistic damage to endothelial cells. Implications for Pseudomonas-associated tissue injury. The Journal of clinical investigation. 1992;90(6):2187–96. Epub 1992/12/01. doi: 10.1172/jci116104. PubMed PMID: 1469082; PubMed Central PMCID: PMCPMC443369.

14. Sorensen RU, Klinger JD. Biological effects of *Pseudomonas aeruginosa* phenazine pigments. Basic Research and Clinical Aspects of Pseudomonas aeruginosa. 1987;39:113–24.

15. Denning GM, Railsback MA, Rasmussen GT, Cox CD, Britigan BE. *Pseudomonas* pyocyanine alters calcium signaling in human airway epithelial cells. American Journal of Physiology-Lung Cellular and Molecular Physiology. 1998;274(6):L893–L900.

16. Lau GW, Hassett DJ, Ran H, Kong F. The role of pyocyanin in *Pseudomonas aeruginosa* infection. Trends in molecular medicine. 2004;10(12):599–606.

17. Hassan HM, Fridovich I. Mechanism of the antibiotic action pyocyanine. Journal of bacteriology. 1980;141(1):156–63. Epub 1980/01/01. doi: 10.1128/jb.141.1.156-163.1980. PubMed PMID: 6243619; PubMed Central PMCID: PMCPMC293551.

18. Yang J, Zhao H-L, Ran L-Y, Li C-Y, Zhang X-Y, Su H-N, et al. Mechanistic insights into elastin degradation by pseudolysin, the major virulence factor of the opportunistic pathogen *Pseudomonas aeruginosa*. Scientific reports. 2015;5(1):1–7.

19. Heck L, Morihara K, Abrahamson D. Degradation of soluble laminin and depletion of tissue-associated basement membrane laminin by *Pseudomonas aeruginosa* elastase and alkaline protease. Infection and immunity. 1986;54(1):149–53.

20. Wilson MJ, McMorran BJ, Lamont IL. Analysis of promoters recognized by PvdS, an extracytoplasmic-function sigma factor protein from *Pseudomonas aeruginosa*. Journal of bacteriology. 2001;183(6):2151–5.

21. Rybtke M, Hultqvist LD, Givskov M, Tolker-Nielsen T. *Pseudomonas aeruginosa* biofilm infections: community structure, antimicrobial tolerance and immune response. Journal of molecular biology. 2015;427(23):3628–45.

22. Olivares E, Badel-Berchoux S, Provot C, Prévost G, Bernardi T, Jehl F. Clinical impact of antibiotics for the treatment of *Pseudomonas aeruginosa* biofilm infections. Frontiers in microbiology. 2020:2894.

23. Moser C, Jensen PØ, Thomsen K, Kolpen M, Rybtke M, Lauland AS, et al. Immune responses to *Pseudomonas aeruginosa* biofilm infections. Frontiers in Immunology. 2021;12:625597.

24. Hassett DJ, Korfhagen TR, Irvin RT, Schurr MJ, Sauer K, Lau GW, et al. *Pseudomonas aeruginosa* biofilm infections in cystic fibrosis: insights into pathogenic processes and treatment strategies. Expert opinion on therapeutic targets. 2010;14(2):117–30.

25. Kuburich NA, Adhikari N, Hadwiger JA. *Acanthamoeba* and *Dictyostelium* Use Different Foraging Strategies. Protist. 2016;167(6):511–25. Epub 2016/10/04. doi: 10.1016/j.protis.2016.08.006. PubMed PMID: 27693864; PubMed Central PMCID: PMCPMC5154823.

26. Erken M, Lutz C, McDougald D. The rise of pathogens: predation as a factor driving the evolution of human pathogens in the environment. Microb Ecol. 2013;65(4):860–8. Epub 2013/01/29. doi: 10.1007/s00248-013-0189-0. PubMed PMID: 23354181; PubMed Central PMCID: PMCPMC3637895.

27. Rayamajhee B, Willcox MD, Henriquez FL, Petsoglou C, Subedi D, Carnt N. *Acanthamoeba*, an environmental phagocyte enhancing survival and transmission of human pathogens. Trends in Parasitology. 2022.

28. Al-Quadan T, Price CT, Abu Kwaik Y. Exploitation of evolutionarily conserved amoeba and mammalian processes by *Legionella*. Trends in microbiology. 2012;20(6):299–306. Epub 2012/04/13. doi: 10.1016/j.tim.2012.03.005. PubMed PMID: 22494803; PubMed Central PMCID: PMCPMC3603140.

29. Kumar A, Molli PR, Pakala SB, Nguyen TMB, Rayala SK, Kumar R. PAK thread from amoeba to mammals. Journal of cellular biochemistry. 2009;107(4):579–85.

30. Siddiqui R, Khan NA. Biology and pathogenesis of *Acanthamoeba*. Parasites & vectors. 2012;5:6. Epub 2012/01/11. doi: 10.1186/1756-3305-5-6. PubMed PMID: 22229971; PubMed Central PMCID: PMCPMC3284432.

31. Segal G, Shuman HA. Legionella pneumophila utilizes the same genes to multiply within *Acanthamoeba castellanii* and human macrophages. Infection and immunity. 1999;67(5):2117–24.

32. Cirillo JD, Cirillo SL, Yan L, Bermudez LE, Falkow S, Tompkins LS. Intracellular growth in *Acanthamoeba castellanii* affects monocyte entry mechanisms and enhances virulence of *Legionella pneumophila*. Infection and immunity. 1999;67(9):4427–34.

33. Moffat JF, Tompkins L. A quantitative model of intracellular growth of *Legionella pneumophila* in *Acanthamoeba castellanii*. Infection and immunity. 1992;60(1):296–301.

34. Khunkitti W, Lloyd D, Furr JR, Russell AD. *Acanthamoeba castellanii*: growth, encystment, excystment and biocide susceptibility. The Journal of infection. 1998;36(1):43–8. Epub 1998/11/20. doi: 10.1016/s0163-4453(98)93054-7. PubMed PMID: 9515667.

35. Kim J-H, Matin A, Shin H-J, Park H, Yoo K-T, Yuan X-Z, et al. Functional roles of mannose-binding protein in the adhesion, cytotoxicity and phagocytosis of *Acanthamoeba castellanii*. Experimental parasitology. 2012;132(2):287–92.

36. Alsam S, Sissons J, Dudley R, Khan NA. Mechanisms associated with *Acanthamoeba castellanii* (T4) phagocytosis. Parasitology research. 2005;96:402–9.

37. Wang X, Ahearn DG. Effect of bacteria on survival and growth of *Acanthamoeba castellanii*. Current microbiology. 1997;34:212–5.

38. Siddiqui R, Lakhundi S, Khan NA. Interactions of *Pseudomonas aeruginosa* and *Corynebacterium* spp. with non-phagocytic brain microvascular endothelial cells and phagocytic *Acanthamoeba castellanii*. Parasitology Research. 2015;114(6):2349–56. doi: 10.1007/s00436-015-4432-0.

39. Abd H, Wretlind B, Saeed A, Idsund E, Hultenby K, Sandström G. *Pseudomonas aeruginosa* utilises its type III secretion system to kill the free-living amoeba *Acanthamoeba castellanii*. Journal of eukaryotic microbiology. 2008;55(3):235–43.

40. Matz C, Kjelleberg S. Off the hook–how bacteria survive protozoan grazing. Trends in microbiology. 2005;13(7):302–7.

41. Shteindel N, Gerchman Y. *Pseudomonas aeruginosa* mobbing-like behavior against *Acanthamoeba castellanii* bacterivore and its rapid control by quorum sensing and environmental cues. Microbiology Spectrum. 2021;9(3):e00642–21.

42. Purdy-Gibson ME, France M, Hundley TC, Eid N, Remold SK. *Pseudomonas aeruginosa* in CF and non-CF homes is found predominantly in drains. J Cyst Fibros. 2015;14(3):341–6. Epub 2014/12/03. doi: 10.1016/j.jcf.2014.10.008. PubMed PMID: 25443472.

43. Remold SK, Brown CK, Farris JE, Hundley TC, Perpich JA, Purdy ME. Differential habitat use and niche partitioning by *Pseudomonas* species in human homes. Microbial ecology. 2011;62:505–17.

44. Purdy-Gibson M, France M, Hundley T, Eid N, Remold S. *Pseudomonas aeruginosa* in CF and non-CF homes is found predominantly in drains. Journal of Cystic Fibrosis. 2015;14(3):341–6.

45. Mojesky AA, Remold SK. Spatial structure maintains diversity of pyocin inhibition in household *Pseudomonas aeruginosa*. Proceedings of the Royal Society B. 2020;287(1938):20201706.

46. Jurado-Martín I, Sainz-Mejías M, McClean S. *Pseudomonas aeruginosa*: An Audacious Pathogen with an Adaptable Arsenal of Virulence Factors. International Journal of Molecular Sciences. 2021;22(6):3128. PubMed PMID: doi:10.3390/ijms22063128.

47. Malone JG. Role of small colony variants in persistence of *Pseudomonas aeruginosa* infections in cystic fibrosis lungs. Infection and drug resistance. 2015:237–47.

48. Hogardt M, Heesemann J. Adaptation of *Pseudomonas aeruginosa* during persistence in the cystic fibrosis lung. International Journal of Medical Microbiology. 2010;300(8):557–62.

49. Sarink M, Pirzadian J, van Cappellen W, Tielens A, Verbon A, Severin J, et al. *Acanthamoeba castellanii* interferes with adequate chlorine disinfection of multidrug-resistant *Pseudomonas aeruginosa*. Journal of Hospital Infection. 2020;106(3):490–4.

50. Hojo F, Osaki T, Yonezawa H, Hanawa T, Kurata S, Kamiya S. *Acanthamoeba castellanii* supports extracellular survival of *Helicobacter pylori* in co-culture. Journal of Infection and Chemotherapy. 2020;26(9):946–54.

51. LaBauve AE, Wargo MJ. Growth and laboratory maintenance of *Pseudomonas aeruginosa*. Current protocols in microbiology. 2012;25(1):6E. 1.-6E. 1.8.

52. Wurtzel O, Yoder-Himes DR, Han K, Dandekar AA, Edelheit S, Greenberg EP, et al. The single-nucleotide resolution transcriptome of Pseudomonas aeruginosa grown in body temperature. 2012.

53. Nielsen MK, Nielsen K, Hjortdal J, Sørensen UBS. Temperature limitation may explain the containment of the trophozoites in the cornea during *Acanthamoeba castellanii* keratitis. Parasitology research. 2014;113:4349–53.

54. Lambrecht E, Baré J, Van Damme I, Bert W, Sabbe K, Houf K. Behavior of *Yersinia enterocolitica* in the presence of the bacterivorous *Acanthamoeba castellanii*. Applied and environmental microbiology. 2013;79(20):6407–13.

55. Baré J, Sabbe K, Huws S, Vercauteren D, Braeckmans K, Van Gremberghe I, et al. Influence of temperature, oxygen and bacterial strain identity on the association of *Campylobacter jejuni* with *Acanthamoeba castellanii*. FEMS microbiology ecology. 2010;74(2):371–81.

56. Sharaby Y, Rodríguez-Martínez S, Pecellin M, Sela R, Peretz A, Höfle MG, et al. Virulence traits of environmental and clinical *Legionella pneumophila* multilocus variable-number tandem-repeat analysis (MLVA) genotypes. Applied and environmental microbiology. 2018;84(10):e00429–18.

