## Supplementary figures and images for "Examining the influence of environmental factors on *Acanthamoeba castellanii* and *Pseudomonas aeruginosa* in co-culture"

### Supplemental Figure 1

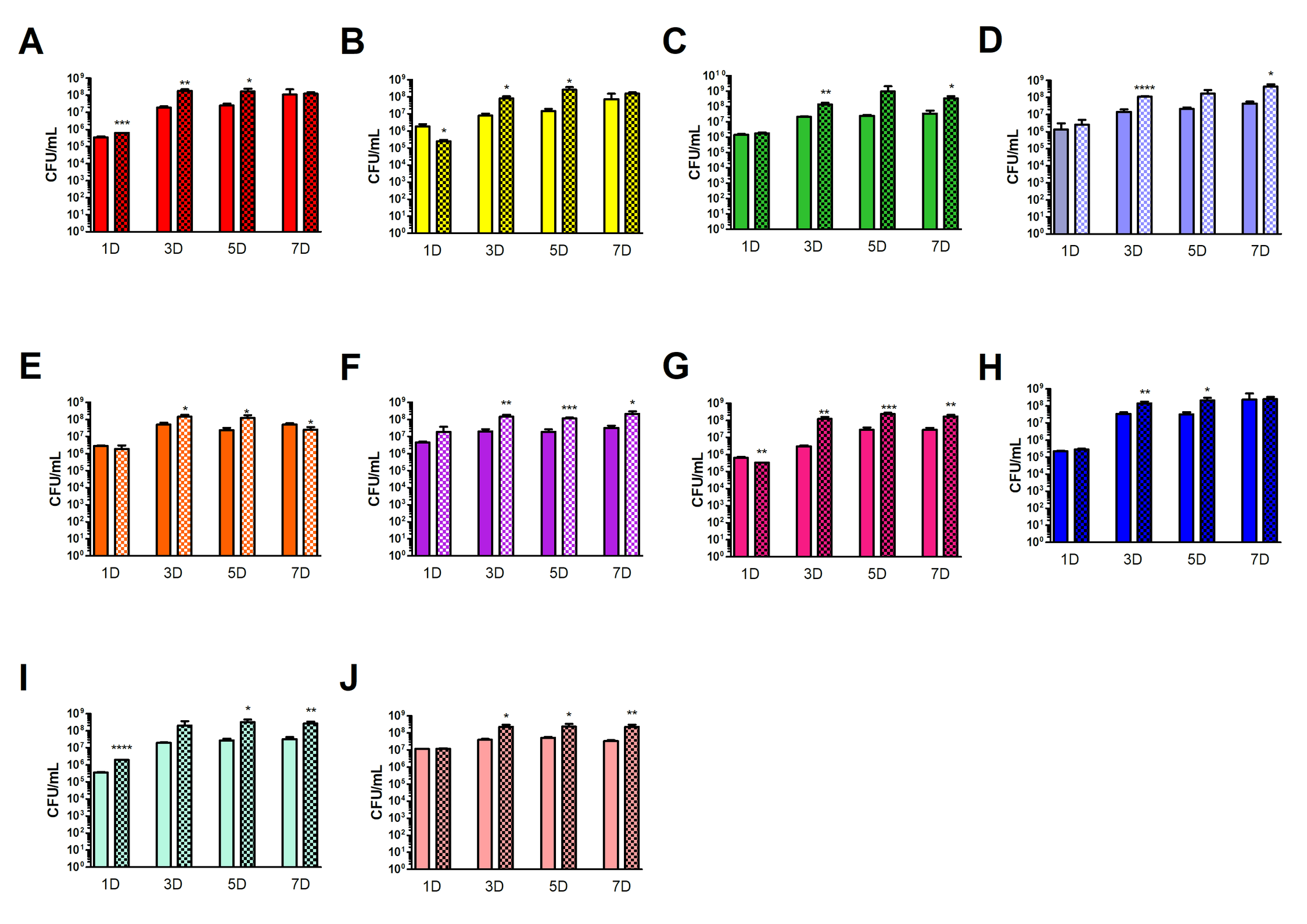

### Supplemental Figure 2

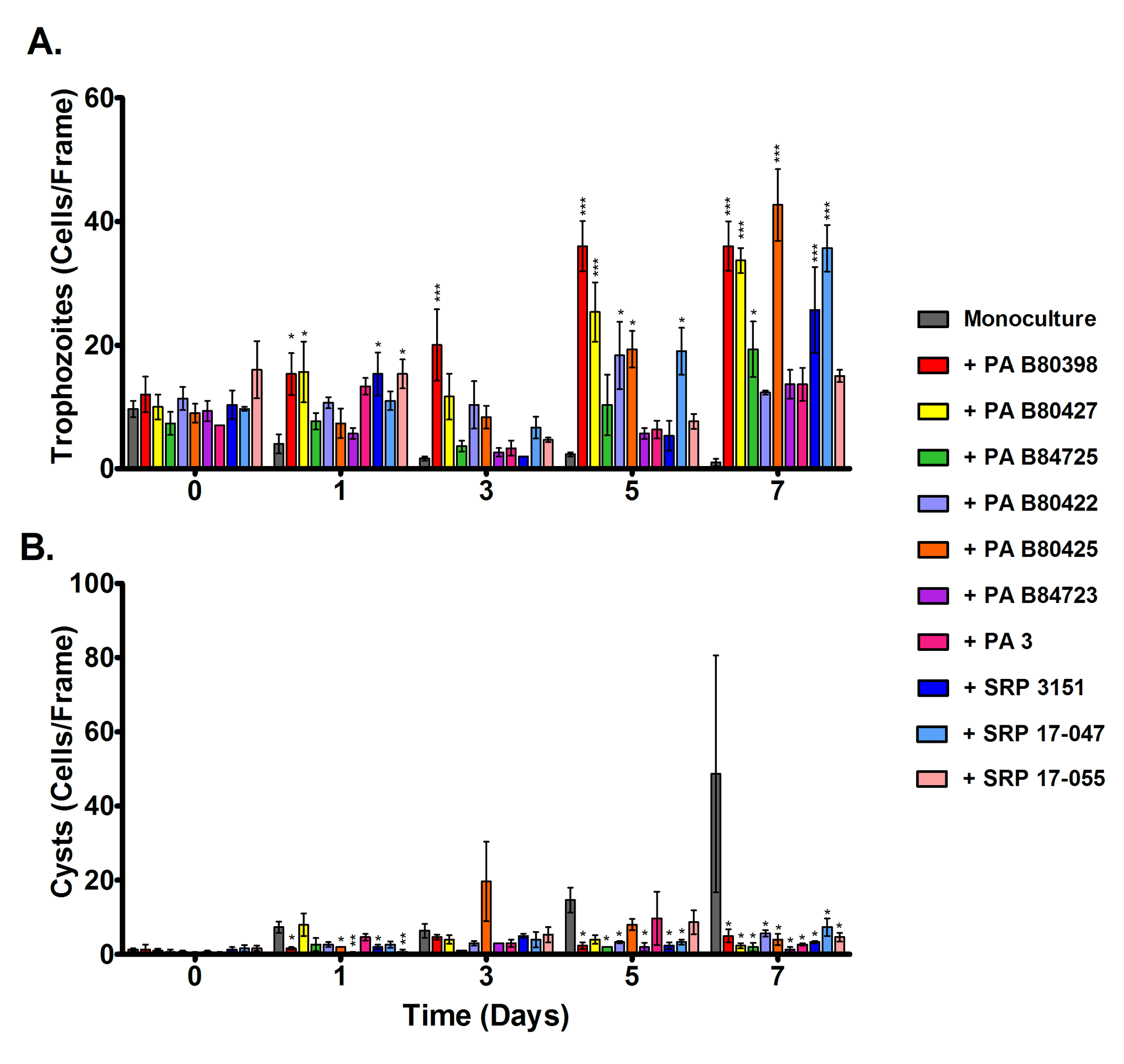
