## Supplemental Table 1 for "Examining the influence of environmental factors on *Acanthamoeba castellanii* and *Pseudomonas aeruginosa* in co-culture"

| | *Pseudomonas aeruginosa* | | | | | *A. castellanii* trophozoites | | | | | *A. castellanii* cysts | | | | |
|---|---|---|---|---|---|---|---|---|---|---|---|---|---|---|---|
| Source of Variation | *Df | *SS | *MS | F | P value | *Df | *SS | *MS | F | P value | *Df | *SS | *MS | F | P value |
| Effect of time (days) | 4 | 2863 | 715.8 | 18.62 | <0.0001 | 4 | 2232 | 558.1 | 62.83 | <0.0001 | 4 | 23.22 | 5.806 | 7.132 | <0.0001 |
| Effect of strain | 9 | 1285 | 142.8 | 3.713 | 0.0005 | 9 | 1225 | 136.1 | 15.32 | <0.0001 | 9 | 10.35 | 1.150 | 1.413 | 0.1926 |
| Interaction (time/strain) | 36 | 4597 | 127.7 | 3.321 | <0.0001 | 36 | 1221 | 33.92 | 3.819 | <0.0001 | 36 | 49.42 | 1.373 | 1.686 | 0.0224 |
| Residual | 100 | 3845 | 38.45 | | | 100 | 888.3 | 8.883 | | | 100 | 81.40 | 0.8140 | | |
